# Development of a genetically-encoded oxytocin sensor

**DOI:** 10.1101/2020.07.14.202598

**Authors:** Neymi Mignocchi, Sarah Krüssel, Kanghoon Jung, Dongmin Lee, Hyung-Bae Kwon

## Abstract

Oxytocin (OXT) is a neuropeptide originating in the paraventricular nucleus (PVN) of the hypothalamus, with a role in influencing various social behaviors. However, pinpointing its actions only during the time animals are performing specific behaviors has been difficult to study. Here we developed an optogenetic gene expression system designed to selectively label neuronal populations activated by OXT in the presence of blue-light, named “OXTR-iTango2”. The OXTR-iTango2 was capable of inducing gene expression of a reporter gene in both human embryonic kidney (HEK) cells and neurons in a quantitative manner. *In vivo* expression of OXTR-iTango2 selectively labeled OXT-sensitive neurons in a blue-light dependent manner. Furthermore, we were able to detect a subset of dopamine (DA) neurons in the ventral tegmental area (VTA) that receive OXT activation during social interaction. Thus, we provide a genetically-encoded, scalable optogenetic toolset to target neural circuits activated by OXT in behaving animals with a high temporal resolution.

## Introduction

Neuromodulation influences animal behavior and perception in space and time. Neuromodulators are incessantly flowing within a brain or body and adjust neurons to react differently when responding to the same inputs and can alter the threshold for synaptic plasticity (S. H. Lee & Dan, 2012; Yagishita et al., 2014). Especially, neuropeptides are small molecules synthesized and released from neurons and affect activity and signaling of neurons. Thus, pinpointing their actions only during the time which animals are performing specific behaviors would be essential to define their roles, but studying neuropeptide functions in behaving animals has not been investigated much.

One of the major difficulties in studying neuropeptide functions *in vivo* is the paucity of appropriate available tools. Previously, neuromodulator actions were monitored in an indirect way. One common way is to monitor neuronal activity by doing Ca^2+^ imaging or electrophysiological recording. We were able to measure the activity pattern changes at individual cells or in a population level. In this way, we may be able to find out significant activity changes mediated by neuromodulators, but it is very difficult to pinpoint where these changes are coming from. Another approach is to directly measure neuromodulator release by fast-scan cyclic voltammetry (FSCV). FSCV can detect endogenous levels of neurotransmitters with high sensitivity (nanomolar scale) and monitor neurotransmitters in behaving animals in real time with a fast temporal resolution (~10 Hz) (Howe, Tierney, Sandberg, Phillips, & Graybiel, 2013; Jennings, 2013). Therefore, quantification is great with this FSCV, but the spatial resolution is poor due to an inserted carbon fiber electrode in the extracellular space of the target brain. Furthermore, FSCV is not able to detect neuropeptides such as oxytocin or vasopressin. More recently, several genetically encoded neuromodulator sensors were developed (Feng et al., 2019; Jing et al., 2018; Patriarchi et al., 2018; Sun et al., 2018). These sensors enable spatiotemporally precise measurements of neuromodulators *in vivo*, but because they are not linked to gene expression, it only visualizes signals, but does not allow the manipulation of target neuron’s functions.

We recently developed a light-gated method to label and manipulate specific neuronal populations activated by neuromodulators in a highly temporally precise manner (Lee et al., 2017). We created an inducible dual protein switch system that is turned on and off by not only a ligand but also light. We named this technique “iTango2”, a light-gated gene expression system that uses β-arrestin and a light-inducible split tobacco-etch-virus (TEV) protease. The iTango2 system allows gene expression by both light and ligand, which means neurons that are activated by neuromodulators can only be labeled when light is illuminated.

In this study, we developed a genetically encoded, scalable OXT receptor iTango2 (OXTR-iTango2). This new optogenetic tool enables selective labeling of OXT sensitive neuronal population with high spatiotemporal resolution. Furthermore, we can quantify the magnitude of OXT release in the target brain areas. We confirmed that OXTR-iTango2 selectively induced gene expression in the presence of OXT and blue light from non-neuronal cells to neurons. Using this sensor, we labeled a population of DA neurons activated by OXT during social interaction. Determining the extent of OXT’s modulation on a specific neuronal population in a mammalian brain will uncover many unknowns of OXT functions.

## Results

### Development of OXTR-iTango2

The molecular design of the iTango2 labeling system allows for the replacement of the expressed GPCRs to label and study other modulatory systems. In this experiment, we cloned the system to contain the OXTR, and then sequenced it to verify the inclusion of the OXTR genetic sequence (Material and methods). OXTR-iTango2 system requires two synthetic proteins and their interaction causes the restoration of split TEV protease functions (Figure 1A). The first protein contains the c-terminal truncated form of OXTR bound to the N-terminus of TEV (TEV-N). The other protein is beta-arrestin2 protein fused with the C-terminus of the split TEV (β-arrestin2-TEV-C). When OXT binds to OXTR, β-arrestin2-TEV-C is bound to OXTR, resulting in the reconstitution of N-TEV and C-TEV protease function. However, in dark condition, TEV protease cannot recognize its substrates because the TEV cleavage sequence is hidden in the improved light-inducible dimer (iLID) (Guntas et al., 2015; D. Lee et al., 2017). iLID is an engineered version of *Avena Sativa* phototropin 1 light-oxygen-voltage 2, (ASLOV2), a protein highly sensitive to blue-light (Guntas et al., 2015). In a dark condition, TEV is unable to access its cleavage site, but upon blue light illumination, iLID conformational changes expose the TEV cleavage sequence, which makes the reconstituted TEV protease recognize the cleavage sequence and release a tetracycline-controlled transcriptional activator (TetR-VP16) out to the cytosol. Then, released TetR-VP16 travels to the nucleus and begins transcription of the reporter gene (Figure 1A).

**Figure 1.**
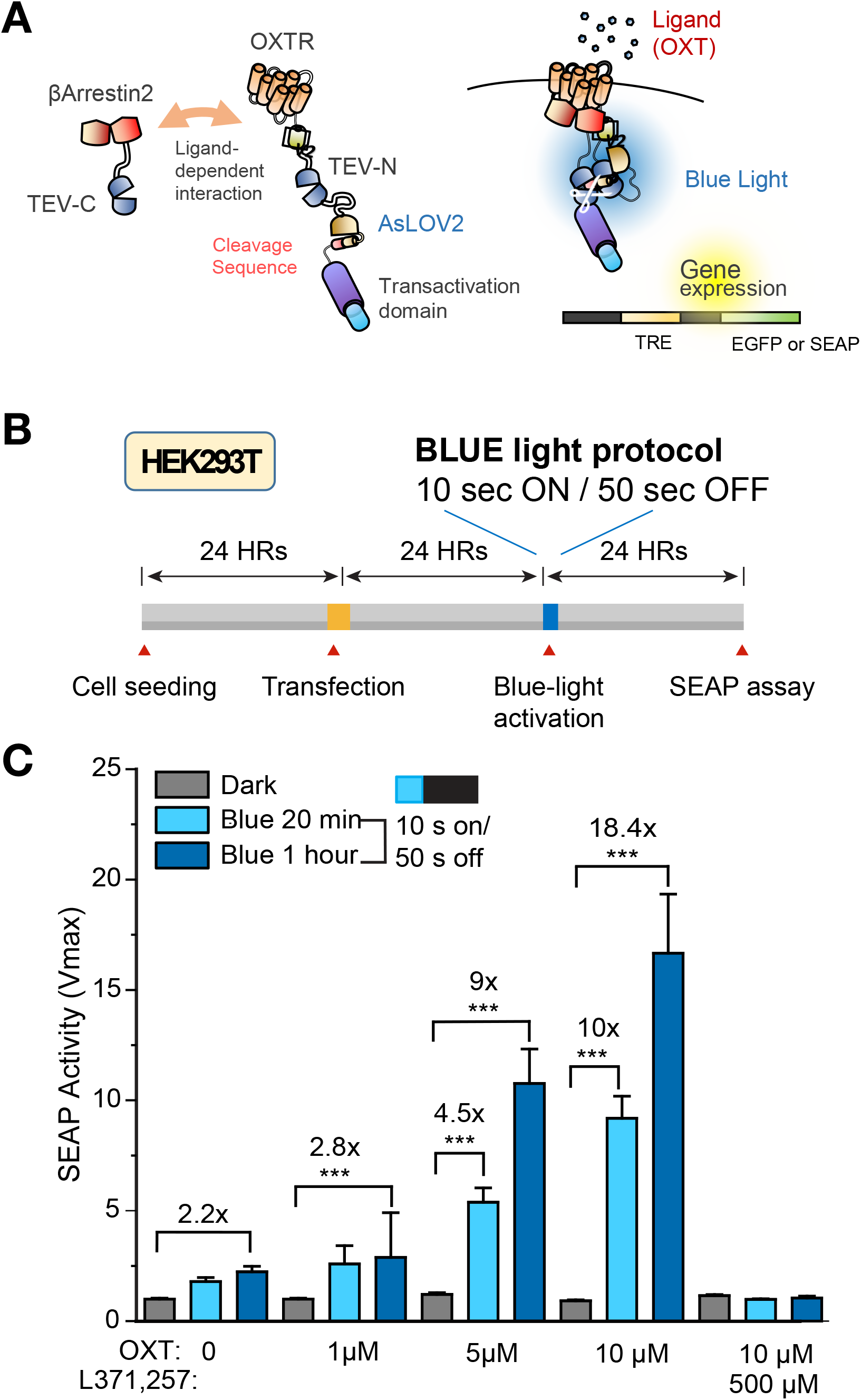
Verification of OXTR-iTango2 in HEK293T cells. **(A)** Graphical illustration of OXTR-iTango2 system. **(B)** Schematic of an experimental plan. **(C)** Summary of OXTR-iTango2 test. Blue light and OXT together lead to robust gene expression in a dose-dependent manner. Longer blue light illumination resulted in higher gene expression. In the absence of light, even high concentration of OXT did not induce gene expression. Gene expression was completely blocked by a selective OXTR antagonist L357,257, indicating resulting gene expression was mediated by OXT. Error bars represent mean ± SEM.

To test the efficiency and reliability of the OXTR-iTango2 system, we expressed the OXTR-iTango2 constructs (i.e. β-arrestin2-TEV-C-P2A-TdTomato and OXTR-V2tail-TEV-N-AsLOV2-tTA) in human embryonic kidney (HEK) 293T cells (Figure 1B). In addition, we co-expressed the secreted alkaline phosphatase (SEAP) gene as a reporter to compare quantitative expression across conditions. Testing of the OXTR-iTango2 gene reporter system included dark and light conditions (i.e. 20 minutes of short blue light pulses of 10 s ON/50 s OFF) in the presence and absence of OXTR agonist OXT. In the dark control, there was no detectable spontaneous gene expression of the SEAP reporter, suggesting that the reporter gene itself does not cause leaky expression. Upon blue light illumination, gene expression was increased by OXT in a dose-dependent manner (Figure 1C). Similarly, the longer blue light was shined, the more genes were expressed. OXT- and light-dependent gene expression was completely blocked by an OXTR antagonist, L371,257, indicating that gene expression was highly selective to OXTR activation (Figure 1C).

### Verification of OXTR-iTango2 in neurons

To test whether OXTR-iTango2 also works in neurons, we generated adeno-associated virus (AAV) expressing OXTR-iTango2 constructs and EGFP reporter genes and infected them into dissociated rat hippocampal neuronal cultures (Figure 2A). Virus-mediated transfection of the OXTR-iTango2 system was applied to the neuronal cultures at 3 days *in vitro* (DIV), and then expressed for 7 days before dividing the cultures into varying blue light conditions. 500 nM concentration of OXT was applied in the absence or the presence of blue light. OXT in dark condition did not cause gene expression significantly, but upon blue light illumination, robust gene expression was made (F(2, 1) = 23.13, p<0.001) (Figure 2B). Although blue light was shined, EGFP expression remained negligible when OXT was not applied to the cultured neurons. Thus, OXTR-iTango2 allows selective labeling of neurons activated by OXT only in the blue light condition with a high signal-to-noise ratio (SNR).

**Figure 2.**
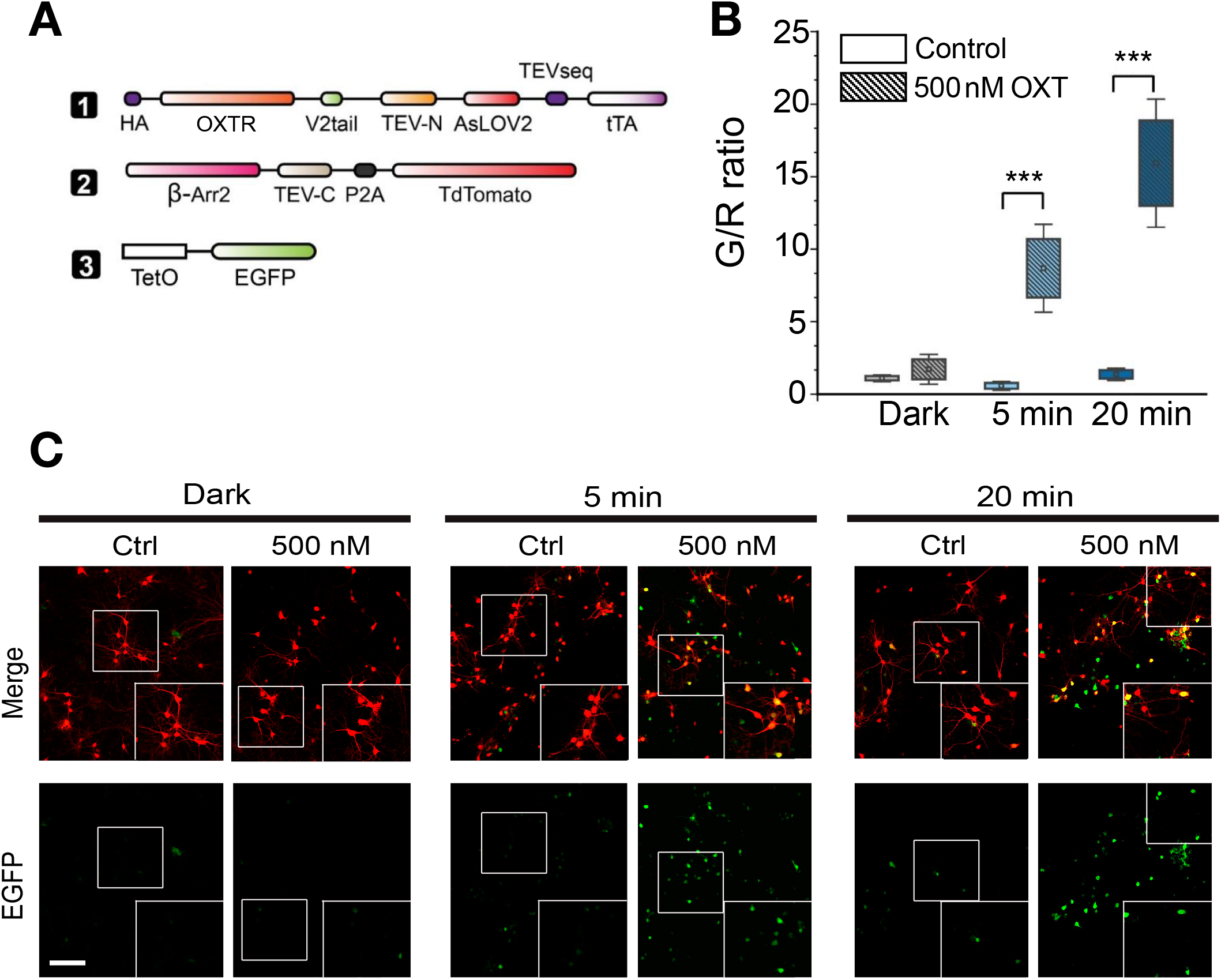
Verification of OXTR-iTango2 in culture neurons. **(A)** A list of viral constructs used for experiments. **(B)** Summary graph demonstrating the expression of EGFP in cultured rat hippocampal neurons in response to different blue-light exposure times and OXT (500nM). The ratio of the intensities of green to red fluorescence (G/R/ratio) was used to measure overall gene expression of OXTR-iTango2 constructs (two-way ANOVA, post hoc Bonferroni test, control versus 500 nM OXT) (Dark/Control,1.09 ± 0.15; Dark/500 nM,1.71 ± 0.69; 5 min/Control, 0.62 ± 0.21; 5 min/500 nM, 8.68 ± 2.02; 20 min/Control, 1.36 ± 0.29; 20 min/500 nM, 15.93 ± 2.00). Squares represent mean and bars represent mean ± SEM. **(C)** Representative confocal images from each condition, showing that the number of EGFP expressing neurons were increased when blue light was illuminated. White boxes (inset) show enlarged images. Scale bars, 200 μm.

### In vivo labeling of an oxytocin-sensitive neuronal population

We further tested the efficacy of OXTR-iTango2 *in vivo.* Previous studies showed that OXT neurons within the paraventricular nucleus (PVN) of the hypothalamus innervate their axons to DA neurons in the ventral tegmental area (VTA), playing a key role in social behaviors (Charlet & Grinevich, 2017; Hung et al., 2017; Tang et al., 2014; Xiao, Priest, & Kozorovitskiy, 2018; Xiao, Priest, Nasenbeny, Lu, & Kozorovitskiy, 2017). We first confirmed the PVN-OXT neuronal axonal projections to VTA-DA neurons through anterograde tracing by expressing AAV1-Flex-EGFP into the PVN of OXT-Cre mice (Figure 3A). Sagittal and coronal brain section images clearly demonstrated direct axonal projections from the PVN-OXT neurons to the VTA-DA neurons (Figure 3B).

**Figure 3.**
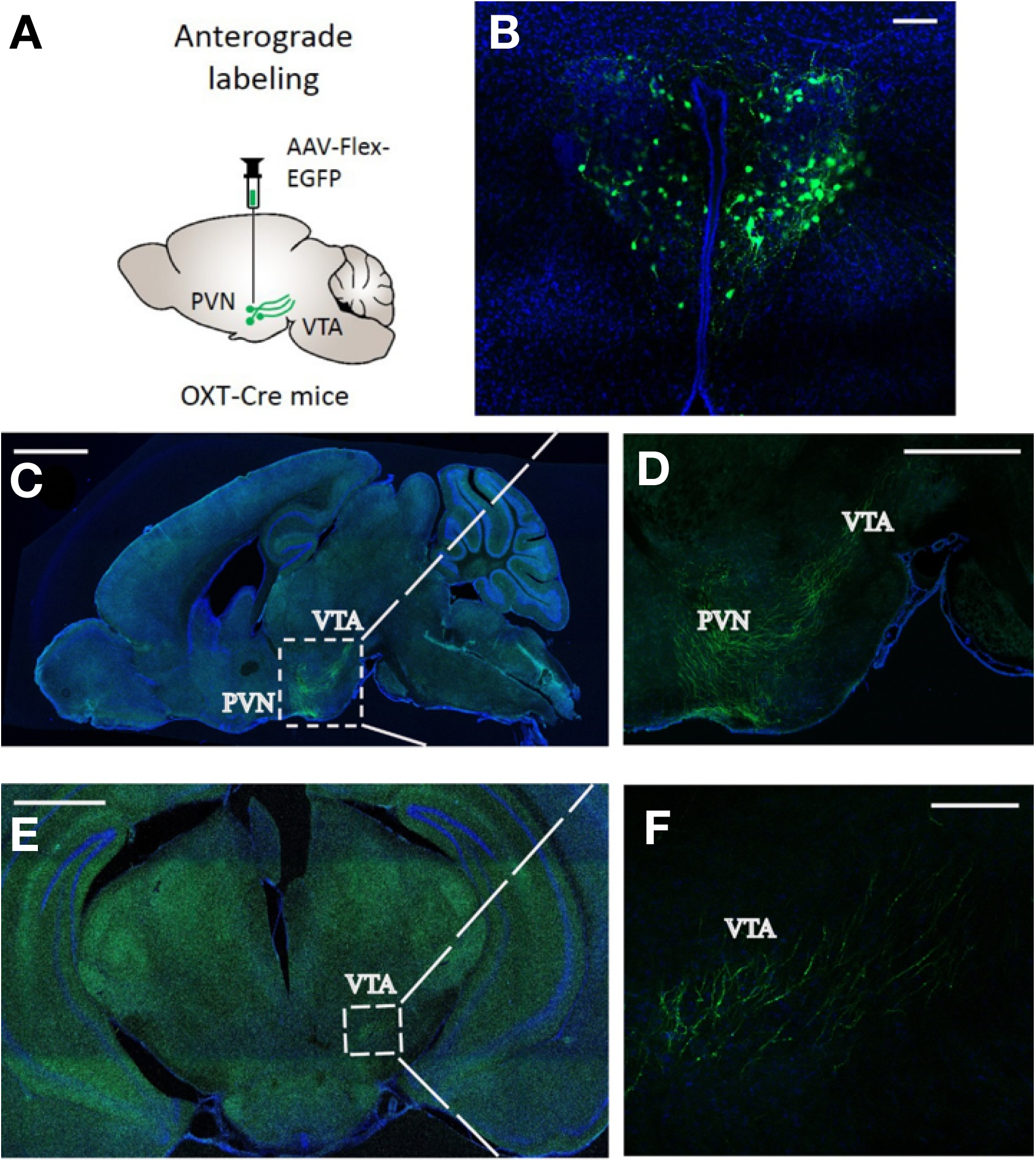
Confirmation of PVN-OXT projections to the VTA. **(A)** Diagram of the AAV1-Flex-EGFP administered into the medial section of the PVN of OXT-Cre mice (top left). Confirmation of EGFP expressing PVN-OXT neurons (top right); Scale bar is 100 μm. **(B)** Sagittal view of the PVN-OXT projections into the VTA (middle left); Scale bar is 2 mm. 10x magnification of sagittal section showing PVN-OXT projections from the PVN to the VTA (middle right); Scale bar is 1 mm. **(C)** A Coronal section view of PVN-OXT in the VTA (bottom left); Scale bar is 1 mm. 10x magnification of coronal section showing PVN-OXT terminals in the VTA (bottom right); Scale bar is 100 μm.

To determine whether OXTR-iTango2 can reliably label neuronal populations *in vivo* with endogenous OXT release, we expressed OXTR-iTango2 viral constructs (AAV1-hSYN-OXTR-V2 tail-TEV-N-AsLOV2-tTA, AAV1-hSYN-β-Arrestin2-TEV-C-P2A-TdTomato, and AAV1-hSYN-TRE-EGFP) into the VTA, and AAV-dFlox-ChR2(H134R)-mCherry (ChR2-mCh) into the PVN of OXT-Cre mice (Figure 4A). Viral injections in both the VTA and PVN were administered in the right hemisphere. We also implanted an optic cannulae to directly shine blue-light in the VTA of mice. In this condition, blue light illumination in the VTA simultaneously triggers OXT release from PVN-OXT terminals and activates OXTR-iTango2-dependent gene expression. As expected, blue light illumination (5 s ON/15 s OFF, for 45 min) significantly induced the expression of the EGFP reporter gene (F_2, 515.6_ = 144.5, p<0.0001) (Figure 4B-4D), with only a marginal expression of EGFP observed in the light and OXT negative control groups (i.e. dark condition and light only condition) (Figure 4B-4D). These results indicate that OXTR-iTango2 constructs do not produce background gene expression of the EGFP reporter gene in the absence of OXT ligand or blue-light. Thus, our results reconfirmed that OXTR-iTango2 is a reliable tool to selectively label OXT sensitive neuronal populations in response to the release of endogenous OXT in the presence of blue-light.

**Figure 4.**
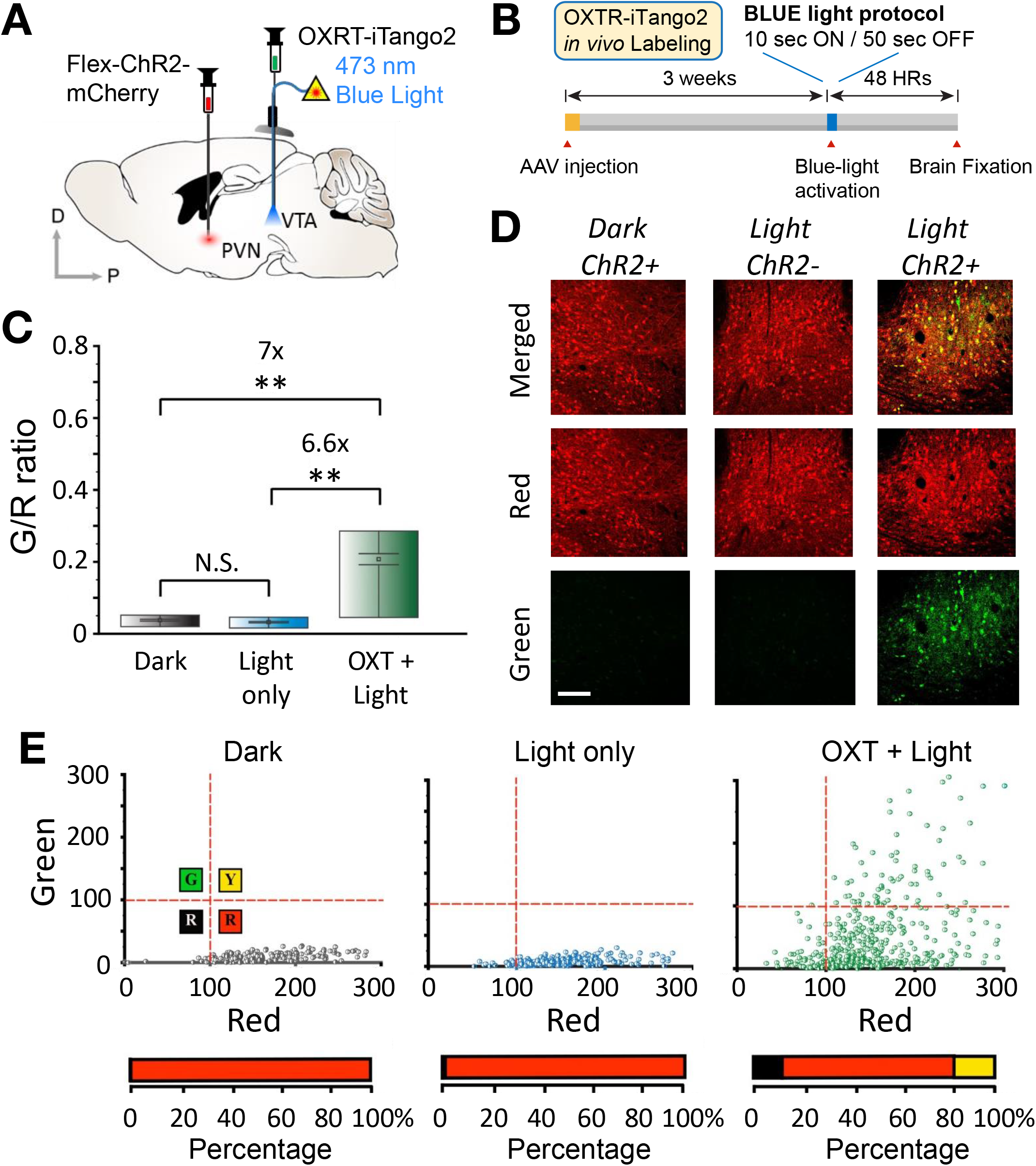
*In vivo* verification of OXTR-iTango2. **(A)** Diagram of the viral injections for *in vivo* verification of OXTR-iTango2. AAV-dFlox-ChR2(H134R)-mCherry was administered into the medial section of the PVN of OXT-Cre mice. A mixture of OXTR-iTango2 viruses including an EGFP reporter were unilaterally injected into the right hemisphere along with an optic fiber to shine blue light and selectively label DA neurons in the transduced hemisphere. **(B)** Summary graph of gene expression *in vivo.* Dark, 3 mice; light only, 3 mice; OXT and light, 3 mice (Welch’s one-way ANOVA, F(2, 515.6) = 144.5, p<0.0001). Boxes represent the 25^th^ and 75^th^ percentiles, and squares represent the mean. Error bars represent mean ± SEM. **(C)** EGFP gene expression in VTA-DA neurons *in vivo.* Blue-light was exposed at a 5 sec ON/15 sec OFF interval for 45 min to release endogenous OXT and expose blue light for EGFP gene expression. Merged (top) and green (bottom) channel images presented. Scale bar, 100 μm. Coronal-section view of high magnification VTA images of a mouse expressing both mCherry and OXTR-iTango2 after blue light exposure. Scale bar 1 mm. **(D)** Gene expression distribution of VTA-DA neuronal populations. Scatter plots (top panel) of two colors (y-axis: EGFP; x-axis: TdTomato) and distribution graph (bottom panel) of VTA cells expressing OXTR-iTango2 across OXT and blue light conditions. Cells analyzed were divided into four quadrants (Green, Yellow, Black, and Red), with their respective percentage concentration within the total population analyzed displayed as horizontal histograms. The red dot crossing lines within the graphs indicates the threshold for defining the quadrants.

### Selective labeling of a subset of neurons during social interaction

OXT is a neuropeptide serving as a neuromodulator in the brain affecting a wide range of social behaviors (Bielsky & Young, 2004; Caldwell, Aulino, Rodriguez, Witchey, & Yaw, 2017; Cavanaugh, Huffman, Harnisch, & French, 2015; de Jong & Neumann, 2018; Feldman, Monakhov, Pratt, & Ebstein, 2016; Love, 2014; M.& A., 2018; Numan & Young, 2016; Sanchez-Andrade & Kendrick, 2009). OXT affects both positive (e.g. trust, maternal behaviors, pair bonding, and partner preference) and negative (e.g. aggression) social interactions and behaviors (Bielsky and Young, 2004; Caldwell et al., 2017; Cavanaugh et al., 2015; de Jong and Neumann, 2018; Feldman, 2017; Gamal-Eltrabily and Manzano-Garcia, 2018; Love, 2014; Numan and Young, 2016; Sanchez-Andrade and Kendrick, 2009). Impaired social interactions skills are a great determinant of various neurophysiological disorders such as autism, depression, and social anxiety disorders (Gunaydin et al., 2014). Manipulating OXT release in the brain affects social interaction behaviors by modulating DA neurons in the VTA (Hung et al., 2017; Xiao et al., 2017). However, it is presumed that OXT was not acting all neurons equally. There should be a neuronal population that are activated by OXT stronger than others during a specific period of behavior. The patterns and the magnitude of OXT activation during social interaction will be important for understanding circuits and algorithms of OXT-mediated behaviors.

To detect OXT-sensitive neurons, we injected viral solution containing OXTR-iTango2 mixed with an EGFP reporter in the VTA. A modified experienced-dependent social conditioning protocol (ESCP) was adopted as a social behavioral task (Hung et al., 2017). In this task, we first assessed if experimental mice have a preferred side by letting them explore freely in the cage (Day 1). Then we put an interacting mouse to the non-preferred side, so we can ensure that social stimuli the experimental mouse receives is mainly caused by another interacting mouse not by internal preference (Figure 5A). On day 2, experimental mice were placed in the middle chamber of a three chamber apparatus, where there was a juvenile mouse in one of the two side chambers, and the other side chamber contained a toy mouse. Experimental mice were conditioned to either a social environment (i.e. juvenile mouse encounter) or a non-social environment (i.e. toy mouse) by allowing them to enter the designated side of the chamber for three 10-minute periods with 5-minute intervals in between each period (the other side chamber remained closed). Each side chamber contained different types of flooring to induce the mouse to associate room environment to the social vs nonsocial environment (Figure 5A). On day 3, we turned on blue light whenever the experimental mouse enters the social chamber side. A whole test session was four 5-min trials with 2-min habituation periods in the middle chamber before each trial. When the mouse stayed in the social side for several minutes, we repeated blue light on/off cycles (3 s ON/12 s OFF) during the stay (Blue + Social). In one control experiment, we performed the same social interaction task, but blue light was not turned on (“Social only”). To ensure that OXT-sensitive neurons are labeled by social interaction, another control experiment was performed where the same blue light procedure was normally applied, when mice entered the toy mouse chamber side (“Blue only”).

**Figure 5.**
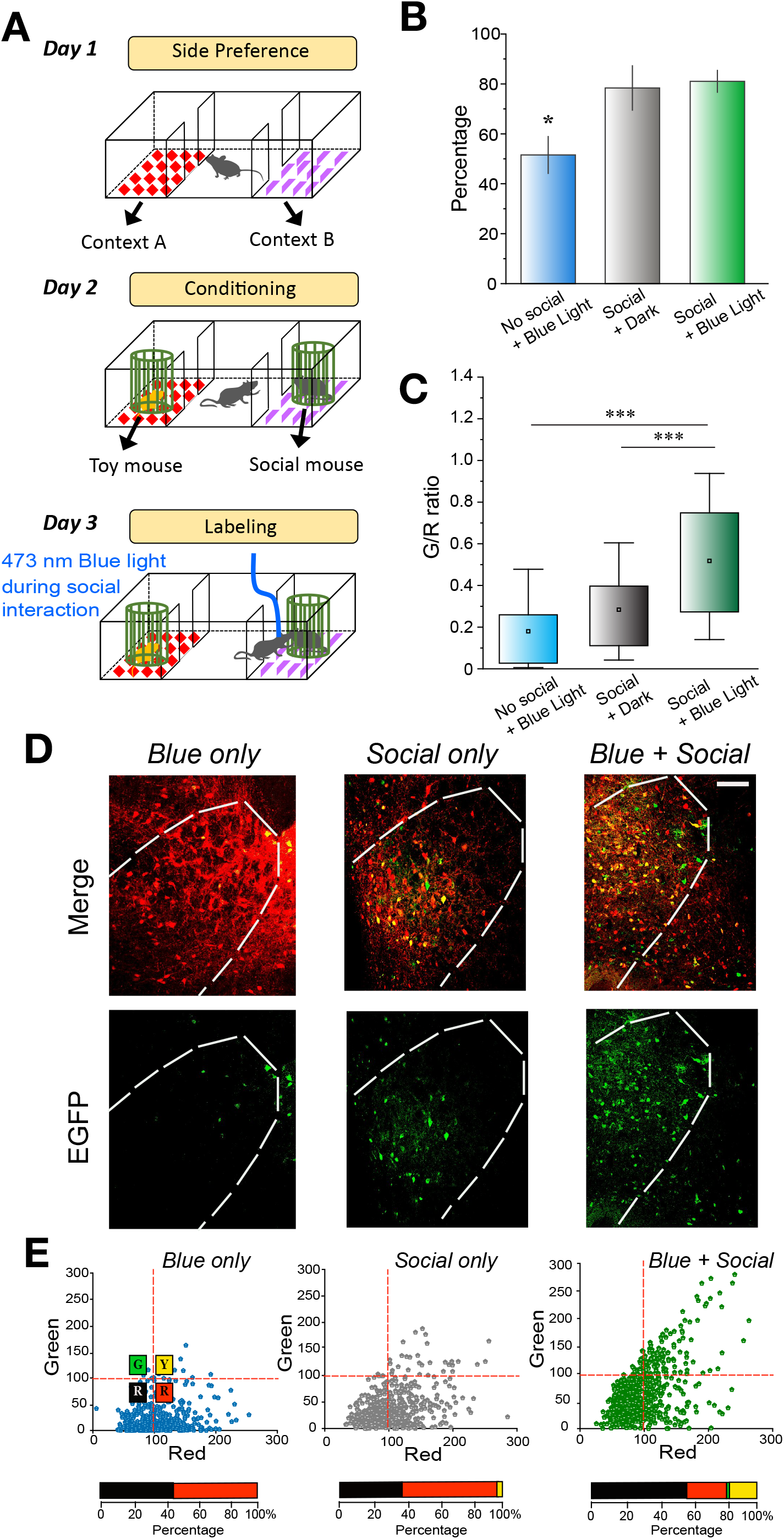
Labeling OXT-sensitive DA neurons during social interaction. **(A)** Diagram of the experienced-dependent social conditioning protocol (ESCP). This protocol was performed to label OXT sensitive VTA-DA neurons during social interactions. On Day 1, mice were exposed to social chamber to determine their preferred floor texture on each side chamber. On Day 2, mice were conditioned to least their preferred side including a social stimulus. On Day 3, testing day, mice receiving blue light exposure upon entry to conditioned or control side chamber. **(B)** Preference to a social environment. Quantification of the percent of time spent from all groups (one-way ANOVA, post-hoc Bonferroni test, F(2,11) = 5.89, p<0.05) (Blue only: Mean = 78.38 ± 8.89, 4 animals; Social only: Mean = 81.05 ± 4.37, 4 animals; Blue + Social: Mean = 51.55 ± 7.37, 6 animals). Error bars represent ± SEM. **(C)** Gene expression of EGFP reporter of OXT-sensitive DA neurons. Summary graph of G/R ratio in the VTA (Blue only = 4 mice; Social only = 4 mice; Blue + Social = 6 mice) (Welch’s one-way ANOVA, F(2,1073) = 249.3, post-hoc Games-Howell, p<0.0001). Boxes show the mean, 25^th^ and 75^th^ percentiles, and whiskers show the 10^th^and 90^th^ percentiles. **(D)** Labeling of VTA-DA neurons with OXTR-iTango2. High magnification representative confocal images of VTA neurons expressing OXTR-iTango2 after blue light exposure. Merged (top) and green (bottom) channel images presented. Scale bar, 100 μm.**(E)** Gene expression distributions of VTA-DA neurons. Scatter plots (top panel) of two colors (y-axis: EGFP; x-axis: TdTomato) and distribution graph (bottom panel) of VTA cells expressing OXTR-iTango2 across social environment conditions. Cells analyzed were divided into four quadrants (Green, Yellow, Black, and Red), with their respective percentage concentration within the total population analyzed displayed as horizontal histograms. The red dot crossing lines within the graphs indicate the threshold for defining the quadrants.

Behavior results showed that mice spent less time in a toy mouse chamber side, suggesting that social experience during the conditioning (Day 2) triggered social reward learning on the next labeling day (Figure 5B). Due to the time difference, we increased the total duration of sessions in the “Blue only” condition, such that total blue light exposure time was matched. We found that EGFP reporter signals were significantly increased in the “Blue + Social” group compared to the “Social only” or the “Blue only” group (Figure 5C). The number of neurons expressing EGFP was also increased in the “Blue + Social” group. These data suggest that higher amount of OXT was released during social interaction and impacted the broader area of the VTA neurons. Thus, OXTR-iTango2 would be a useful technique that can quantify the OXT release in live animals and capture the specific ensemble of neurons related to OXT-mediated functions in space and time.

## Discussion

The hormone OXT plays a central role in the development of motivated social behaviors including mate bonding and preference, mother/offspring bonding, aggression and sexual behavior (Bielsky & Young, 2004; Cavanaugh et al., 2015; Feldman & Bakermans-Kranenburg, 2017; Love, 2014; M.& A., 2018; Numan & Young, 2016; Sanchez-Andrade & Kendrick, 2009). The vast concentration and distribution of OXTR throughout the mesocorticolimbic DA pathway (Peris et al., 2017; Shahrokh, Zhang, Diorio, Gratton, & Meaney, 2010) suggests that OXT has a role in influencing rewarding and motivated social behaviors.

In this study, we aim to develop a genetically encoded, scalable OXT sensor that can visualize neuronal population activated by OXT in the OXT-DA pathways involved in social rewards in response to learned social stimuli. This new optogenetic tool would enable selective labeling of OXT sensitive neuronal population in high spatiotemporal resolution. Furthermore, developing a way to detect OXT-sensitive neurons will be beneficial for many investigators who want to use this technique in their studies.

Initially, we screened our newly cloned OXTR-iTango2 in HEK293T cell cultures and confirmed gene expression of a reporter gene is dependent on the presence of OXT and blue-light exposure. The next set of experiments was performed in neural cultures to induce the expression of an EGFP reporter gene under blue light and OXT conditions with results showing OXTR’s efficacy in translating OXT and blue-light dependent conditions into significantly increases of EGFP reporter expression. After this successful labeling of neuronal cultures, we set to test OXTR-iTango2 *in vivo* once we confirmed OXT-neural projections (PVN-OXT) onto VTA-DA neurons, and determined its ability to induce gene expression with endogenous OXT. Our *in vivo* experiments provided visual evidence of the reliability of OXTR-iTango2 to label OXT-sensitive cell populations (VTA-DA neurons) with a high spatial and temporal resolution in the combined presence of endogenous OXT and blue-light.

Altogether, our *in vitro* and *in vivo* experiments examining the efficacy and reliability of this newly developed OXT-sensor, OXTR-iTango2, suggests that it is a promising tool that can be used to dissect neural circuits involved in OXT-influenced mammalian behaviors.

## Materials and Methods

### Construction of OXTR-iTango2

To generate CMV::HA-OXTR-V2Tail-TEN-V-iLID1-TetR, we first PCR amplified the OXTR sequence (Addgene, #66467) to contain ClaI restriction enzyme sites at each end. The amplified OXTR product and CMV-DRD2-iTango2 construct (D. Lee et al., 2017) were both digested with the ClaI restriction enzyme. Then, we ligated the digested OXTR-PCR product into the iTango2 backbone to generate the final CMV-OXTR-iTango2 DNA product. OXTR-iTango2 was then sequenced (Eurofin Genomics) to confirm the presence of OXTR within the construct. All reagents and enzymes used for cloning were purchased from New England Biolabs.

### HEK293T cell culture preparation, DNA transfections, and blue-light exposure

Human Embryonic Kidney (HEK) 293T cells were acquired from the American Type Culture Collection (ATCC). These cells were chosen for initial testing of OXTR-iTango2 because they do not possess neuronal OXTRs or channels, which means that they maintain an environment to detect effects caused by transfected neuronal proteins. Large batches of HEK293T cells were grown in high-glucose Dulbecco’s Modified Eagle’s Medium (DMEM) (Gibco) with 1% penicillin-streptomycin (Invitrogen) and 10% FBS (Gibco, 10438-018), while maintained in incubated conditions at 37°C and 10% CO2. Experiments required the preparation of 24-well plates coated with a 1mg/ml poly-D-lysine hydrobromide (Sigma-Aldrich, P0899) solution for two hours and washed off with distilled water before cell plating. Batches of HEK293T cells were then treated with 0.25% trypsin (Gibco, 25200) for 2 minutes to detach them from the surface of the plates used to grow them. The detached cells were collected and then counted to plate a total of 2.0 X 10^5^ cells per 12-mm PDL coated well. 24 hours after plating, a Calcium Phosphate Transfection kit (Clontech) was used to transfect cells with a mixture of OXTR-iTango2 DNA plasmid vectors with a secreted alkaline phosphatase (SEAP) enzyme reporter gene (4:1:4 ratio; OXTR-TEV-N-iLID1-tTA, β-Arrestin2-TEV-C, TRE-SEAP). The OXTR-iTango2 plasmid vectors were first slowly added into 2x HEPES-buffered saline and incubated for 1 hour to precipitate. The precipitated solution was then added into each well in equal amounts. On the third day, HEK293T cells received various concentrations of OXT and/or L371,257 (OXTR antagonist) before exposing them to blue-light illumination with a 465-nm-wavelength blue LED array (LED Wholesalers) at an interval time of 10 s ON/50 s OFF. Blue-light interval was controlled automatically by a high-accuracy digital electronic timer (GraLab, model 451). The LED array was located inside of an incubator set to 37°C and 10% CO_2_, and we placed a clear and empty plate with a height of 2-cm between the LED light source and the cell cultures. We used a power meter (Thorlabs, Inc, PM100D) to measure the power of the blue-light and set it to 1.7 mW. The dark condition was achieved by covering the plate with aluminum foil. Media changes were performed under a dim red lighting. 48 hours after blue-light exposure, we measured for gene expression of the reporter SEAP enzyme.

### SEAP chemoluminescence assay

We used the secreted embryonic alkaline phosphatase (SEAP) chemoluminescent assay to quantify gene expression by OXTR-iTango2 in HEK293T cells. 40 μL medium samples from each cell-containing well in the various OXT concentrations and blue-light conditions were collected and placed in 96-well plates. Samples were placed inside of a 50-60°C incubator for 10 minutes to inhibit the activity of endogenous alkaline phosphatase before adding a mix of SEAP substrates and L-homoarginine. We set a microplate reader (Molecular Devices, SpectraMax Plus 384) to 37°C and measured the chemoluminescence of the samples at 405 nm every 30 seconds for a total of 2 hours. The reagents used to run the assay were acquired from InvivoGen. We used SoftMax Pro 5.4.1 (Molecular Devices) to measure and calculate Vmax calculations for all samples.

### Dissociated rat hippocampal neuron culture preparation, DNA transfections, and blue-light exposure

Primary dissociated neuronal cultures were performed as described in previous literature (D. Lee et al., 2012). To briefly describe the process, CD IGS rat hippocampus (embryonic day 18-19) were quickly dissected and digested in 0.25% trypsin-EDTA (Invitrogen) at 37°C for 8-10 minutes. Trypsin-EDTA was then removed, and the hippocampal brain tissue was gently triturated ~10-15 times using a 100-1,000 μL pipette tip. 12-mm PDL-coated coverslips (Neuvitro) were placed on 24-well plates, and dissociated cells were counted to divide and add 10^5^ cells to each coverslip containing well. The medium used to plate neurons consisted of neurobasal medium (Invitrogen) with 1% FBS (Thermo Fisher Scientific), 1% Glutamax supplement (Gibco), and 2% B27 supplement (Gibco). Every 3-4 days, one-half of the media was replaced with freshly prepared medium lacking FBS. On DIV 3, cultures were infected with diluted (1:10) OXTR-iTango2 viral constructs (1:1:2 ratio; AAV1-hSYN-OXTR-TEV-C-P2A-iLiD-tTA, AAV1-hSYN-β-Arrestin2-TEV-N-P2A-TdTomato, AAV1-TRE-EGFP). A week later (DIV 10), varying concentrations of OXT (Tocris, Cat #: 1910) and/or L-371,257 (Tocris, Cat #: 2410) were added to the neuronal cultures and illuminated with blue light using the same time intervals and equipment described above for varying periods of time (5 min, 20 min, and 60 min). After the light protocol was completed, the medium was completely replaced with new medium to remove OXT and/or L-371,257. On DIV 12, all neurons were fixed with 4% paraformaldehyde for imaging.

### Animals

OXT-IRES-Cre (OXT-Cre) heterozygous mice (6-10 weeks old, Jackson Laboratory, stock #:024234) and C57BL/6J (Jackson Laboratory, Cat #:664) were used for experiments. Control and test group animals were randomly chosen for social behavior experiments. All of the experimental protocols complied with the National Institute of Health guidelines, and approved by the Max Planck of Florida Institute for Neuroscience Institutional Animal Care and Use Committee and the Johns Hopkins University Animal Care and Use Committee.

### Adeno-associated viral constructs

AAV1-hSYN-OXTR-TEV-C-P2A-iLiD-tTA plasmid was first cloned in the laboratory and then sent to ViGene Bioscience for virus production. All other viral constructs were directly purchased or produced from the ViGene Bioscience. The list of viruses are as follows: (1) AAV_1_-hSYN-OXTR-TEV-C-P2A-iLiD-tTA, (2) AAV1-hSYN-β-Arrestin2-TEV-C-P2A-TdTomato (Addgene Cat #: 89873), (3) AAV1-TRE-EGFP (Addgene Cat #: 89875), (4) AAV1-dFlox-ChR2(H134R)-mCherry (Addgene Cat #: 20297).

### Animal surgeries and stereotactic viral injections

Mouse brains were injected with AAV viral solutions using a stereotactic setup (Kopf instruments). Mice were first fully anesthetized with a ketamine (80 mg/Kg) and xylazine (12.5mg/kg) (Sigma-Aldrich) cocktail, administered through an i.p. injection. Petroleum ointment (Puralube Vet Ophthalmic Ointment) was administered to both eyes to maintain moisture and prevent drying during surgery. Before performing a craniotomy, each mouse had the hair on its head (i.e. surgical region) removed using hair-remover lotion (Nair, Church & Dwight). Mice’s head was then fixed to the stereotactic setup with the use of an ear bar and a nose clamp. The surgical region was prepped with 10% betadine solution (Purdue Product LP). The mouse’s body temperature was maintained at 37°C using a homeothermic blanket placed beneath the surgical space that contained a flexible probe (Harvard Apparatus) during the entirety of the surgery. Before proceeding, animals were checked to determine if they were fully anesthetized. We then carefully removed the skin on the head and periosteum with the use of sharp scissors and scalpel respectively, all within aseptic surgical conditions. The bregma and lambda lines were used to guide and adjust maintenance of brain area location. We used a handheld drill (Fordom Electric Co.) to make a small burr hole (~0.5 mm in diameter) in the skull where the pipette containing premixed viral injections were applied. The iTango2 viral injections consisted of 1:1:2 ratio; AAV1-hSYN-OXTR-TEV-C-P2A-iLiD-tTA, AAV1-hSYN-β-Arrestin2-TEV-N-P2A-TdTomato, AAV1-TRE-EGFP with a 500 nl injection volume; and dFlox-ChR2(H134R)-mCherry viral solution (500 nl). The anterograde tracing of OXT neuron projection from the PVN to the VTA was performed by injecting 400 nl of flex-EGFP into the PVN of OXT-Cre mice. Viral injections were applied through microinjections with a steady flow and controlled rate of ~0.400-0.500 nl/min using a constructed glass micropipette (Braubrand, tip size 10-20 μm diameter) that was connected to a syringe pump device (World Precision Instruments). Glass pipettes used for microinjections were constructed using a micropipette puller (P-1000, Sutter Instruments), and the tip of each micropipette was angled and sharpened using a micropipette grinder (EG-400, Narishige). We used the Mouse Brain Atlas and preliminary injection trials with fast green dye to determine the injection coordinates of brain areas of interest. The following coordinates were used for each brain area: (1) VTA, AP −3.2 mm, ML +1.25 mm from bregma, DV −4.35 mm from the brain’s surface at a 10° angle; and (2) PVN, AP −0.75 mm, ML +1.5 mm from bregma, DV 4.9 mm from the brain’s surface at a 15° angle. After the entirety of the premixed virus was injected into the brain, the tip of the micropipette was held in place for 3 minutes before removal to prevent any backflow of viral solutions.

### Fabrication and implantation of optic fibers

Optic fibers (Thorlabs, low OH, 200-μm core, 0.37 NA; BFL37-2000) were first cut with a diamond knife and then inserted into a 1.25-mm-diameter ceramic ferrule (Thorlabs, CFLC230-10) using a 230-μm bore. We used epoxy (Gorilla Glue Company) to bond the optic fiber to the ferrule. Once bonded, polishing sandpaper and a grinding puck (Thorlabs) were used to finely grind both ends of the optic fiber. After each viral injection, the optic fiber was implanted perpendicularly into the targeted brain area using a cannulae holder (Thorlabs, XCL), and the aid of the stereotactic device. To ensure the security of the optic fiber, dental cement (Parkell, C&B-Metabond) was applied to adhere the optic fiber to the skull of each animal. Once the dental cement was dried to its entirety, we removed the cannulae holder from the implanted and secured optic fiber. An analgesic (buprenorphine, 0.05 mg kg^-1^ body weight) was injected subcutaneously to each animal upon the completion of each surgery to alleviate any postsurgical pain before returning mice to their home cages for surgical recovery.

### In vivo blue-light labeling

After surgery, mice were allowed to recover from virus injections and optic-fiber implantations for 17-19 days before optogenetic manipulation was performed. Blue-light at ~10mW power was applied with the use of a 473-nm laser (Changchun New Industries Optogenetic Technology, MBL-FN-473). Blue-light was applied in a 5 s ON/15 s OFF interval for a total of 45 minutes through the implanted optic fiber (Thorlabs, 200-μm core, 0.37 NA). We controlled the timing of the light delivery using MATLAB (Mathworks) to generate a custom code.

### Experience-dependent social conditioning protocol (ESCP) training and labeling using OXTR-iTango2

The ESCP protocol was adapted from Hung et al. (2017). After animals underwent surgery for viral injections and optic-fiber implantation, they were allowed to recover for 17-22 days before undergoing behavior training and OXTR-iTango2 dependent labeling. One week before the beginning of testing, mice were placed in individual cages and cages were placed in a reverse-cycle room to ensure activity and motivation during the trial days.

On day 1 (D1) of training, pre-conditioned animals were placed in a 3-social chamber with two different flooring textures in each side chamber (sandpaper vs a metal wire sheet) to determine if they had a preferred texture. D1 was performed to decrease any intrinsic differences. Animals were allowed to freely move in the chamber for 20 minutes until they reached a 65-70% preference to one texture. After establishing their baseline preference between the textures of each side chamber, we socially conditioned the animals to the side chamber with their least preferred texture on day 2 (D2). D2 conditioning was composed of a 2-minute habituation period before opening their assigned side chamber and allowing animals 10 minutes to explore the chamber now containing a social stimulus (juvenile mouse, P21-35). The habituation periods involved the animals being enclosed in the middle chamber without the ability to see through the walls to each side chamber. The conditioning protocol was repeated for a total of 3 times with the purpose of conditioning mice to the environment (i.e. floor texture) that contained a rewarding social stimulus. On day 3 (D3), animals underwent testing (i.e. familiar social exposure or a toy mouse) and blue-light labeling of OXT sensitive VTA-DA neurons. Testing was composed of four-5 min trials with 2-minute habituation periods in the middle chamber before each trial. Animals were given access to their now conditioned side after each habituation period was completed and blue-light illumination was applied for 3 seconds immediately upon entry of side chamber (i.e. encounter with conditioned stimulus, floor texture) at ~13-15mW power using a 473-nm laser (Changchun New Industries Optogenetic Technology, MBL-FN-473). Optogenetic stimulation was applied every time the mouse entered the side chamber during the each 5-minute trial session or 3 s ON/12 s OFF when the mouse was staying in the side chamber for longer than 15 seconds. In order to match the total duration of blue light exposure between “Blue only” and “Blue + Social” groups, trial sessions were repeated more for the “Blue only” group. The onset of the optogenetic stimulation was controlled by monitoring each animal’s behaviors in real-time through a Noldus Ethovision camera. 48 hours after blue-light illumination, mice underwent transcardial perfusions after deeply anesthetizing them with a ketamine and xylazine mixture to preserve and dissect brains for confocal imaging (Zeiss LSM880).

### Immunocytochemistry

EGFP and TdTomato staining for neuronal cultures was performed as follows: (1) individual fixed cultures were rinsed three times in PBS, pH 7.4; (2) cultures were blocked in 10% normal donkey serum (Jackson ImmunoResearch, 017-000-121) and 0.1% Triton-X for 30 minutes and inserted into a shaking incubator set to 23°C at 120-130RMP; (3) cultures were incubated with a mixture of RFP antibody pre-absorbed (1:1000 in blocking reagent, Rockland antibodies & assays, 600-401-379) and GFP antibody (1:1000 in blocking reagent, Abcam, ab13970) for 90 minutes in a shaking incubator at room; (4) cultures were rinsed in PBS three times with 5-minute incubation periods each time; (5) cultures were incubated in Cy3-AffiniPure Donkey Anti-Rabbit IgG (H+L) (1:500, Jackson ImmunoResearch, 711-165-152) and Alexa Fluor 488 AffiniPure Donkey Anti-Chicken IgG (H+L) (1:500, Jackson ImmunoResearch, 703-545-155) for 30 minutes in a shaking incubator set to room temperature; (6) cultures were again rinsed with PBS three times with 5-minute incubation periods. All cultures were imaged with a confocal microscope (Zeiss LSM880).

### Statistics

The statistical significance of the SEAP expression in HEK293T cell cultures was calculated using a two-way ANOVA post-hoc Bonferroni test in Origin 2020 software. The statistical significance of the G/R of neuronal cell cultures was also calculated using a two-way ANOVA with post-hoc Bonferroni test in Origin 2020 software. Comparison of G/R labeling *in vivo* was performed with a nonparametric Welch’s one-way ANOVA Test. Single, double, and triple significance asterisks present *P<0.05, **P<0.01, and ***P<0.005.

## Acknowledgements

We thank members of the Kwon laboratory for helpful discussions. This work was supported by Johns Hopkins University School of Medicine (to H-B.K.), Max Planck Florida Institute for Neuroscience (to H-B.K.), the National Institutes of Health Grants R01MH107460 (to H-B.K.), and DP1MH119428 (to H-B.K).

## References

Bielsky, I. F., & Young, L. J. (2004). Oxytocin, vasopressin, and social recognition in mammals. Peptides, 25(9), 1565–1574. https://doi.org/10.1016/j.peptides.2004.05.019

Bosch, O. J., Nair, H. P., Ahern, T. H., Neumann, I. D., & Young, L. J. (2009). The CRF system mediates increased passive stress-coping behavior following the loss of a bonded partner in a monogamous rodent. Neuropsychopharmacology. https://doi.org/10.1038/npp.2008.154

Buffington, S. A., Di Prisco, G. V., Auchtung, T. A., Ajami, N. J., Petrosino, J. F., & Costa-Mattioli, M. (2016). Microbial Reconstitution Reverses Maternal Diet-Induced Social and Synaptic Deficits in Offspring. Cell, 165(7), 1762–1775. https://doi.org/10.1016/j.cell.2016.06.001

Caldwell, H. K., Aulino, E. A., Rodriguez, K. M., Witchey, S. K., & Yaw, A. M. (2017). Social context, stress, neuropsychiatric disorders, and the vasopressin 1b receptor. Frontiers in Neuroscience, 11(OCT), 1–10. https://doi.org/10.3389/fnins.2017.00567

Cavanaugh, J., Huffman, M. C., Harnisch, A. M., & French, J. A. (2015). Marmosets treated with oxytocin are more socially attractive to their long-term mate. Frontiers in Behavioral Neuroscience, 9(OCT), 1–11. https://doi.org/10.3389/fnbeh.2015.00251

Charlet, A., & Grinevich, V. (2017). Oxytocin Mobilizes Midbrain Dopamine toward Sociality. Neuron. https://doi.org/10.1016/j.neuron.2017.07.002

de Jong, T. R., & Neumann, I. D. (2018). Oxytocin and aggression. In Current Topics in Behavioral Neurosciences. https://doi.org/10.1007/7854_2017_13

Dölen, G., Darvishzadeh, A., Huang, K. W., & Malenka, R. C. (2013). Social reward requires coordinated activity of nucleus accumbens oxytocin and serotonin. Nature, 501(7466), 179–184. https://doi.org/10.1038/nature12518

Ebert, A., & Brüne, M. (2018). Oxytocin and social cognition. In Current Topics in Behavioral Neurosciences. https://doi.org/10.1007/7854_2017_21

Feldman, R., & Bakermans-Kranenburg, M. J. (2017). Oxytocin: a parenting hormone. Current Opinion in Psychology, 15, 13–18. https://doi.org/10.1016/j.copsyc.2017.02.011

Feldman, R., Monakhov, M., Pratt, M., & Ebstein, R. P. (2016). Oxytocin Pathway Genes: Evolutionary Ancient System Impacting on Human Affiliation, Sociality, and Psychopathology. Biological Psychiatry, 79(3), 174–184. https://doi.org/10.1016/j.biopsych.2015.08.008

Feng, J., Zhang, C., Lischinsky, J. E., Jing, M., Zhou, J., Wang, H., … Li, Y. (2019). A Genetically Encoded Fluorescent Sensor for Rapid and Specific In Vivo Detection of Norepinephrine. Neuron, 102(4), 745–761.e8. https://doi.org/10.1016/j.neuron.2019.02.037

Gunaydin, L. A., Grosenick, L., Finkelstein, J. C., Kauvar, I. V., Fenno, L. E., Adhikari, A., … Deisseroth, K. (2014). Natural neural projection dynamics underlying social behavior. Cell, 157(7), 1535–1551. https://doi.org/10.1016/j.cell.2014.05.017

Guntas, G., Hallett, R. A., Zimmerman, S. P., Williams, T., Yumerefendi, H., Bear, J. E., & Kuhlman, B. (2015). Engineering an improved light-induced dimer (iLID) for controlling the localization and activity of signaling proteins. Proceedings of the National Academy of Sciences of the United States of America, 112(1), 112–117. https://doi.org/10.1073/pnas.1417910112

Howe, M. W., Tierney, P. L., Sandberg, S. G., Phillips, P. E. M., & Graybiel, A. M. (2013). Prolonged dopamine signalling in striatum signals proximity and value of distant rewards. Nature, 500(7464), 575–579. https://doi.org/10.1038/nature12475

Hung, L. W., Neuner, S., Polepalli, J. S., Beier, K. T., Wright, M., Walsh, J. J., … Malenka, R. C. (2017). Gating of social reward by oxytocin in the ventral tegmental area. Science, 357(6358), 1406–1411. https://doi.org/10.1126/science.aan4994

Jennings, K. A. (2013). A comparison of the subsecond dynamics of neurotransmission of dopamine and serotonin. ACS Chemical Neuroscience, 4(5), 704–714. https://doi.org/10.1021/cn4000605

Jing, M., Zhang, P., Wang, G., Feng, J., Mesik, L., Zeng, J., … Li, Y. (2018). A genetically encoded fluorescent acetylcholine indicator for in vitro and in vivo studies. Nature Biotechnology, 36(8). https://doi.org/10.1038/nbt.4184

Lee, D., Creed, M., Jung, K., Stefanelli, T., Wendler, D. J., Oh, W. C., … Kwon, H. B. (2017). Temporally precise labeling and control of neuromodulatory circuits in the mammalian brain. Nature Methods, 14(5), 495–503. https://doi.org/10.1038/nmeth.4234

Lee, D., Lee, H. W., Hong, S., Choi, B. Il, Kim, H. W., Han, S. B., … Kim, H. (2012). Inositol 1,4,5-trisphosphate 3-kinase A is a novel microtubule-associated protein: PKA-dependent phosphoregulation of microtubule binding affinity. Journal of Biological Chemistry, 287(19), 15981–15995. https://doi.org/10.1074/jbc.M112.344101

Lee, S. H., & Dan, Y. (2012). Neuromodulation of Brain States. Neuron, 76(1), 209–222. https://doi.org/10.1016/j.neuron.2012.09.012

Liu, X., Liu, S., Huang, R., Chen, X., Xie, Y., Ma, R., … Zhang, X. (2018). Neuroimaging studies reveal the subtle difference among social network size measurements and shed light on new directions. Frontiers in Neuroscience, 12(JUL), 1–6. https://doi.org/10.3389/fnins.2018.00461

Love, T. M. (2014). Oxytocin, motivation and the role of dopamine. Pharmacology Biochemistry and Behavior. https://doi.org/10.1016/j.pbb.2013.06.011

M., G.-E., & A., M.-G. (2018). Role of central oxytocin and dopamine systems in nociception and their possible interactions: Suggested hypotheses. Reviews in the Neurosciences. https://doi.org/10.1515/revneuro-2017-0068 LK - http://sfx.library.uu.nl/utrecht?sid=EMBASE&issn=03341763&id=doi:10.1515%2Frevneuro-2017-0068&atitle=Role+of+central+oxytocin+and+dopamine+systems+in+nociception+and+their+possible+interactions%3A+Suggested+hypotheses&stitle=Rev.+Neurosci.&title=Reviews+in+the+Neurosciences&volume=29&issue=4&spage=377&epage=386&aulast=Gamal-Eltrabily&aufirst=Mohammed&auinit=M.&aufull=Gamal-Eltrabily+M.&coden=RNEUE&isbn=&pages=377-386&date=2018&auinit1=M&auinitm=

Meyer-Lindenberg, A., Domes, G., Kirsch, P., & Heinrichs, M. (2011). Oxytocin and vasopressin in the human brain: Social neuropeptides for translational medicine. Nature Reviews Neuroscience, 12(9), 524–538. https://doi.org/10.1038/nrn3044

Numan, M., & Young, L. J. (2016). Neural mechanisms of mother-infant bonding and pair bonding: Similarities, differences, and broader implications. Hormones and Behavior. https://doi.org/10.1016/j.yhbeh.2015.05.015

Parker, K. J., Oztan, O., Libove, R. A., Sumiyoshi, R. D., Jackson, L. P., Karhson, D. S., … Hardan, A. Y. (2017). Intranasal oxytocin treatment for social deficits and biomarkers of response in children with autism. Proceedings of the National Academy of Sciences of the United States of America, 114(30), 8119–8124. https://doi.org/10.1073/pnas.1705521114

Patriarchi, T., Cho, J. R., Merten, K., Howe, M. W., Marley, A., Xiong, W. H., … Tian, L. (2018). Ultrafast neuronal imaging of dopamine dynamics with designed genetically encoded sensors. Science, 360(6396). https://doi.org/10.1126/science.aat4422

Peris, J., MacFadyen, K., Smith, J. A., de Kloet, A. D., Wang, L., & Krause, E. G. (2017). Oxytocin receptors are expressed on dopamine and glutamate neurons in the mouse ventral tegmental area that project to nucleus accumbens and other mesolimbic targets. Journal of Comparative Neurology. https://doi.org/10.1002/cne.24116

Sanchez-Andrade, G., & Kendrick, K. M. (2009). The main olfactory system and social learning in mammals. Behavioural Brain Research, 200(2), 323–335. https://doi.org/10.1016/j.bbr.2008.12.021

Shahrokh, D. K., Zhang, T. Y., Diorio, J., Gratton, A., & Meaney, M. J. (2010). Oxytocin-dopamine interactions mediate variations in maternal behavior in the rat. Endocrinology, 151(5), 2276–2286. https://doi.org/10.1210/en.2009-1271

Sun, F., Zeng, J., Jing, M., Zhou, J., Feng, J., Owen, S. F., … Li, Y. (2018). A Genetically Encoded Fluorescent Sensor Enables Rapid and Specific Detection of Dopamine in Flies, Fish, and Mice. Cell, 174(2), 481–496.e19. https://doi.org/10.1016/j.cell.2018.06.042

Tang, Y., Chen, Z., Tao, H., Li, C., Zhang, X., Tang, A., & Liu, Y. (2014). Oxytocin activation of neurons in ventral tegmental area and interfascicular nucleus of mouse midbrain. Neuropharmacology, 77, 277–284. https://doi.org/10.1016/j.neuropharm.2013.10.004

Xiao, L., Priest, M. F., & Kozorovitskiy, Y. (2018). Oxytocin functions as a spatiotemporal filter for excitatory synaptic inputs to VTA dopamine neurons. ELife, 7, 1–26. https://doi.org/10.7554/eLife.33892

Xiao, L., Priest, M. F., Nasenbeny, J., Lu, T., & Kozorovitskiy, Y. (2017). Biased Oxytocinergic Modulation of Midbrain Dopamine Systems. Neuron, 95(2), 368–384.e5. https://doi.org/10.1016/j.neuron.2017.06.003

Yagishita, S., Hayashi-Takagi, A., Ellis-Davies, G. C. R., Urakubo, H., Ishii, S., & Kasai, H. (2014). A critical time window for dopamine actions on the structural plasticity of dendritic spines. Science, 345(6204), 1616–1620. https://doi.org/10.1126/science.1255514

